# Sampling requirements and approaches to detect ecosystem shifts

**DOI:** 10.1101/2020.04.07.030643

**Authors:** Rosalie Bruel, Easton R. White

## Abstract

Environmental monitoring is a key component of understanding and managing ecosystems. Given that most monitoring efforts are still expensive and time-consuming, it is essential that monitoring programs are designed to be efficient and effective. In many situations, the expensive part of monitoring is not sample collection, but instead sample processing, which leads to only a subset of the samples being processed. For example, sediment or ice cores can be quickly obtained in the field, but they require weeks or months of processing in a laboratory setting. Standard sub-sampling approaches often involve equally-spaced sampling. We use simulations to show how many samples, and which types of sampling approaches, are the most effective in detecting ecosystem change. We test these ideas with a case study of Cladocera community assemblage reconstructed from a sediment core. We demonstrate that standard approaches to sample processing are less efficient than an iterative approach. For our case study, using an optimal sampling approach would have resulted in savings of 195 person-hours—thousands of dollars in labor costs. We also show that, compared with these standard approaches, fewer samples are typically needed to achieve high statistical power. We explain how our approach can be applied to monitoring programs that rely on video records, eDNA, remote sensing, and other common tools that allow re-sampling.

## Introduction

Environmental monitoring is one of the core components to modern ecosystem research and management (McDonald-Madden et al. 2010; White 2019; Lindenmayer et al. 2020). Within an adaptive management framework, monitoring is needed for both learning about the system under study and assessing the effectiveness of management interventions (Lovett et al. 2007). Increasingly, long-term monitoring programs, like the Long Term Ecological Research (LTER) Network in the USA, are becoming available (Maguran et al. 2010). However, environmental monitoring is still often expensive and time-consuming, especially when further processing is needed following sample collection (Zhang and Zhang 2012). Thus, for many fields there is a disparity between the amount of data that can be acquired and stored, and the ultimate number of samples that can be processed. Therefore, monitoring programs need to be designed in such a way to address the question of interest while using limited resources efficiently (Legg and Nagy 2006; McDonald-Madden et al. 2010; Lindenmayer et al. 2020).

Accounting for key considerations in the design of a monitoring program is necessary to detect long-term ecological change. The specifics of the monitoring program will determine the power with which a question of interest can be addressed. For example, White (2019) found that 72% of vertebrate populations required at least 10 years of monitoring to detect significant changes in the population size over time. The specific number of years required depended on the species biology and the detection method used (White 2019). Other work has focused on the frequency of monitoring (Wauchope et al. 2019), showing that if a significant trend is observed from a dataset with limited temporal perspective, it is likely to reliably describe qualitatively the complete trend (increase or decline), but is unlikely to provide an accurate quantitative change in population. Other research has investigated the impact of allocating monitoring resources spatially versus temporally (Rhodes and Jonzen 2011; Weiser et al. 2019), and the benefit of increasing sampling breadth relying on citizen science (Weiser et al. 2020). Lastly, both the ecological and economical costs of failing to detect a true trend (type II error) have to be weighed against the risks of false (type I error) detection (Mapstone 1995). Given limited budgets, monitoring programs need to be designed to be cost-effective (Caughlan 2001; Grantham et al. 2008; Bennett, Rühland, and Smol 2016).

Because of new technological advances, there are many data sources that can be derived long after the actual processes occurred. For example, sediment cores can be retrieved from aquatic ecosystems with little sediment disturbances, such as lakes or lagoon, allowing reconstruction of past ecological communities or conditions (Cohen 2003). Similarly, environmental samples (e.g. water, soil) can be saved and processed later for composition, including eDNA (Bohmann et al. 2014). Likewise, photo- or video-based monitoring can record snapshots of a system and be analyzed later (O’Connell, Nichols, and Karanth 2011; Mallet and Pelletier 2014). In each of these cases, decisions have to made about how much data to extract from the previously collected samples (Zhang and Zhang 2012). Should the paleoecological core be analyzed at every centimeter? Should the video be assessed once per minute? As long as processing samples is expensive, these trade-offs will remain.

Here, we develop a set of tools to determine the appropriate number of samples and sampling approach when dealing with data sources where only a subset of samples are analyzed. We tailor our analysis to the detection of a changepoint in a time series, but our approach is applicable to other questions as well. Changepoints are an important characteristic of a time series as they can indicate a underlying change in ecosystem processes (James and Matteson 2014). We focus on paleoecological core samples as one example of this type of data. We examine the situation where the goal is to detect the time at which a significant change in an ecological community occurs, i.e. a changepoint. However, our approaches are widely applicable to other questions and data types. We first investigate these tools using a simulation-based approach. We then test the tools on a case study from a paleosequence of Cladocera community assemblage from Lake Varese located in the subalpine region of north-western Italy (Bruel et al. 2018).

## Sampling approaches and changepoint detection

For both our simulations and case study, we investigate the effect of different sampling strategies on our ability to detect a changepoint. We begin by either creating simulated time series or using an actual paleoecological time series (Fig. 1). We then subsampled each time series (White and Bahlai 2020) to test the effect of three different sampling approaches along with varying the sample size (Fig. 1c-f). We compared the estimated changepoint from the subsampled time series to that of the full time series as a measure of the effectiveness.

**Figure 1:**
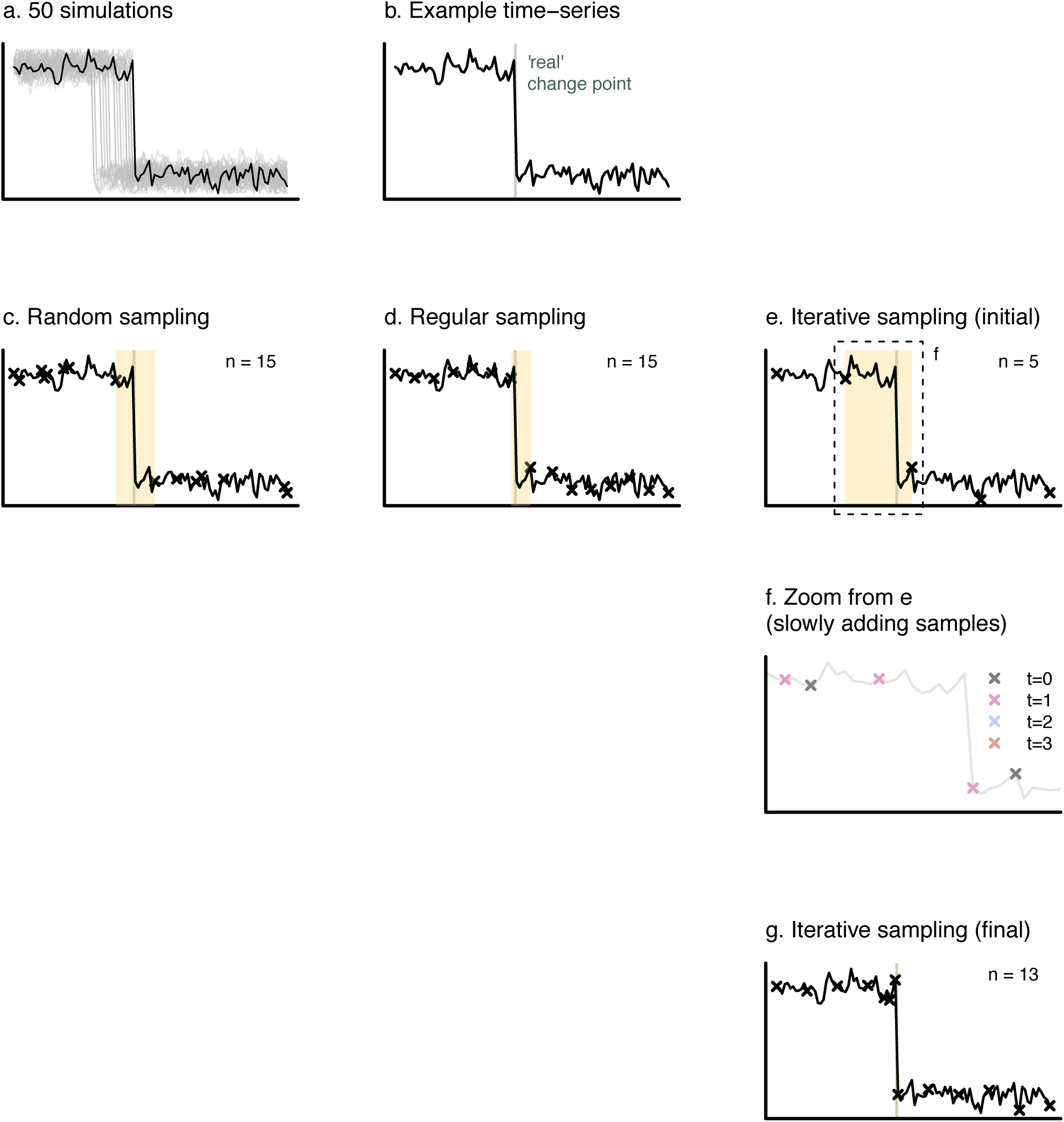
Conceptual diagram illustrating the process of taking (a) simulations of a time series and (b) selecting a single simulation to analyze with three different sampling approaches: (c) random, (d) regular, and (e) iterative. The iterative sampling approach requires (f) adding samples around a detected changepoint until (g) a certain level of accuracy is achieved.

The random sampling approach involves taking a set number of random points throughout the time series (Fig. 1c). In the context of sediment cores, this would mean analyzing community composition at random locations along the core. Random sampling is recommended in designs aimed at quantifying the average size of a population (spatial approach) (Nad’o and Kaňuch 2018). We hypothesize that random sampling will perform the worst in estimating the changepoint. Regular sampling is commonly used (e.g., pigments in Milan et al. (2015)) and requires that a set number of samples be taken at regular intervals (Fig. 1d). Lastly, iterative sampling involves first taking a set number of samples and then iteratively adding samples until a pre-determined level of precision is achieved (Fig. 1e-g). For each scenario, we begin by sampling the first and last sample to ensure coverage of the whole time period. We describe each approach in more detail in the supplementary material and provide code.

We detect changepoints with the function *e.divisive* in the R package *ecp* (James, Zhang, and Matteson 2019). There are several methods available to detect changepoint (reviewed in James and Matteson (2014)); *e.divisive* is a divisive hierarchical estimation algorithm for multiple change point analysis. We chose this method because it is able to perform multiple change point analysis for both uni- and multi-variate time series, without a priori knowledge of the number of changepoints. Herein, we focus on detecting the most important changepoint (i.e. the one of largest magnitude), although we tested the method on a time-series that would have multiple changepoints (Fig. S3). In order to test the performance, we detected the “true” changepoint on the whole time-series, and compare the changepoint found on the sub-sample with the “true” one. The distance to true changepoint served as the performance diagnostic.

## Simulation approach

### Simulation model

We began with a theoretical exploration of the sampling requirements to detect a changepoint. We modeled a simple first order autoregressive (AR-1) process (the discrete-time version of the Ornstein–Uhlenbeck process) with a response variable (*X*_*t*_) that represents either population size, biodiversity, or some other unidimensional metric of community composition at time *t*. The model includes temporal autocorrelation (*ϕ*), the mean of the process (*μ*_*X*_), and a white noise term (*w*_*t*_). The white noise term is a normal distribution with mean (*μ*_*w*_) and variance (*σ*^2^):

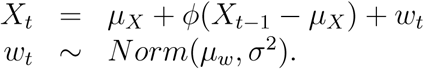

We included a changepoint by shifting *μ*_*X*_ at time *τ* given a specific shift size (*δ*).

We explored how each of these model parameters affected our ability to detect a change point. We simulate an entire time series to serve as the “true” data for comparison (White and Bahlai 2020). We specifically study how the number of samples and the type of sampling affects the detection probability. For simulations, statistical power is the fraction of simulations that were able to detect a changepoint. We define an accurate changepont detection if an estimate is within five time points (given a time series of 100 time points) of the true changepoint. The minimum number of samples required is the number needed for 0.8 statistical power.

## Simulation results

In line with theory on optimal monitoring, we found that the probability of correcting identifying a changepoint decreased with smaller levels of population variability (*σ*) and autocorrelation (*ϕ*) (Fig. 2). We also found that the probability of correct changepoint detection increases with larger shift sizes, which is essentially the effect size (Fig. 2). There were interaction effects between the variables. For example, autocorrelation was only important if population variability was high (Fig. 2). Thus, the number of samples required to obtain high statistical power (above 0.8) increased with larger population variability, lower autocorrelation, and smaller shift sizes (Fig. 3). As predicted, iterative sampling performed best, followed by regular and random sampling (Fig. 3). The distance to the true changepoint, and consequently the minimum number of samples required, was lower for iterative sampling. (Fig. 3)

**Figure 2:**
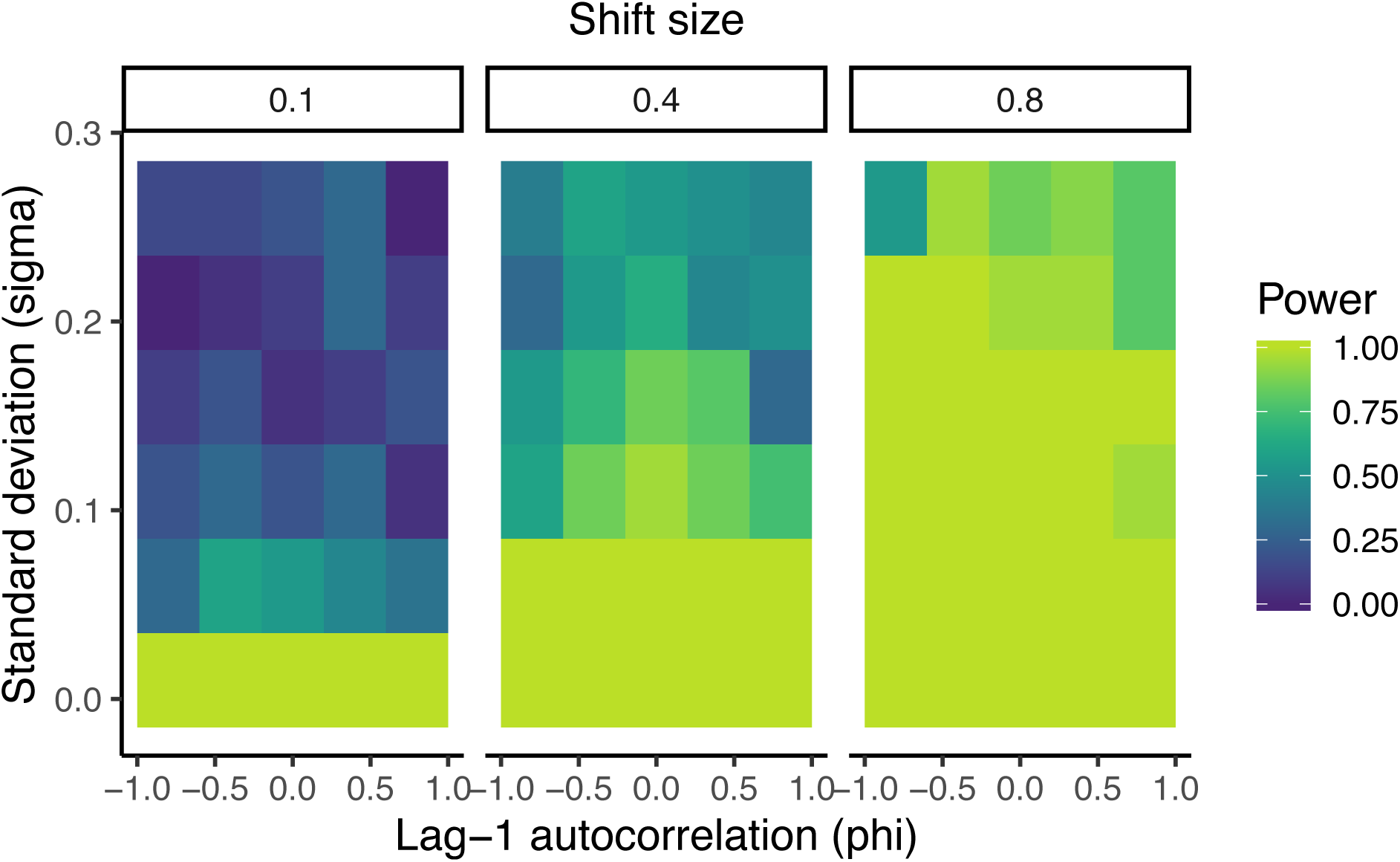
Regular sampling statistical power (fraction of 100 simulations which detected a changepoint within five time points of the true changepoint) for different levels of standard deviation (*σ*), lag-1 autocorrelation (*ϕ*), and shift size (*δ*). For each parameter combination, 20 samples were used. An increase in samples would increase the statistical power across this graph.

**Figure 3:**
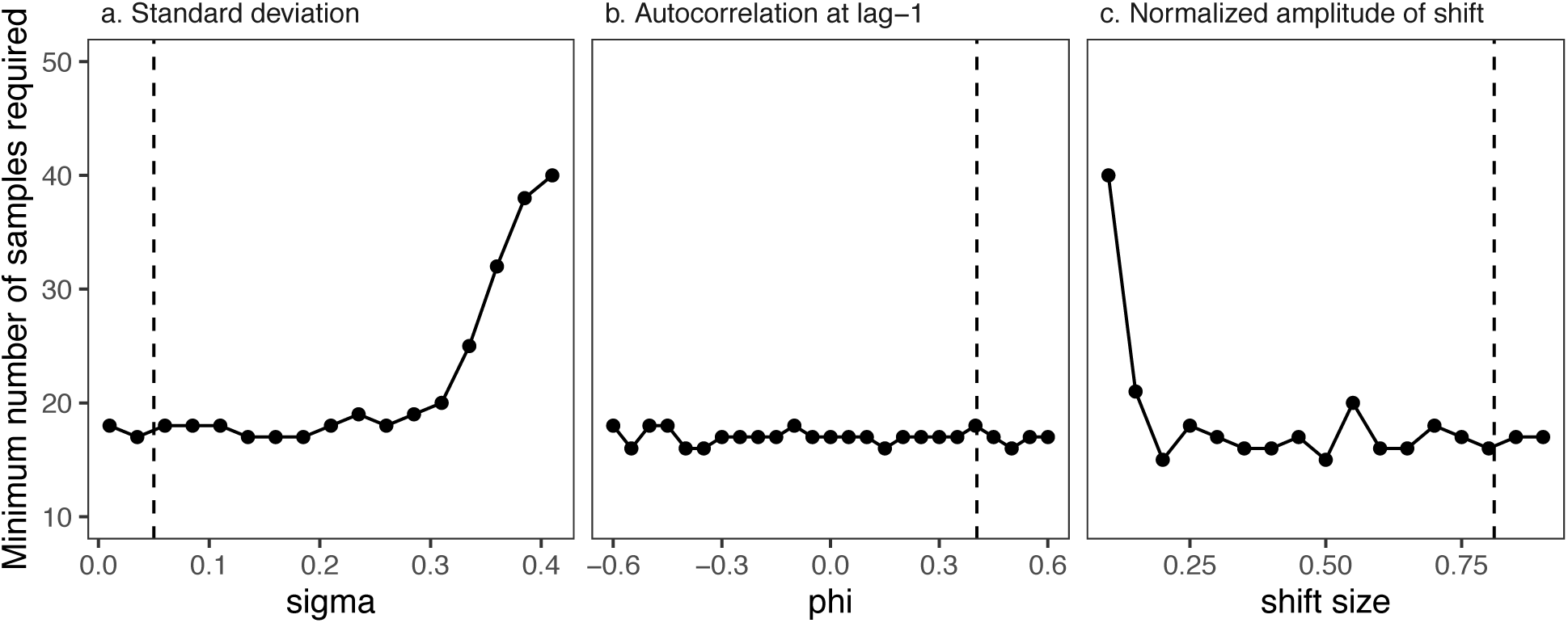
Minimum number of samples required for 0.8 statistical power given different levels of (a) temporal variability (*σ*), (b) temporal autocorrelation (*ϕ*), and (c) shift size (*δ*). Regular sampling was used with the default parameters: *σ* = 0.53, *ϕ* = 0.404, and a shift size = 0.81 to match the case study. The exact timing of the true changepoint varied for each simulation between time steps 30 and 70. Vertical line indicate respective parameter calculated from the case study time series. The same analyses for random and iterative sampling is in Figs. S1,S2.

## Case study

### Case study background

We examined the performance of our approach to detect changepoints in a paleosequence. Paleolimnology provides a reconstruction of past environments over long periods of time, under the premise that sedimentation was not perturbed (low mixing and disturbances). A sediment core is typically subsampled to narrow down periods of time to be compared, either at regular intervals (e.g., Milan et al. (2015)), or continuously (e.g., Perga et al. (2015)).

We tested different sampling methods on a real community time series from Lake Varese (IT), with the objective to detect the main changepoint in Cladocera assemblage. Lake Varese is a small (14.8 km^2^), deep (z_*max*_= 26m), monomictic lake, in the subalpine region of north-western Italy (238 m asl). It underwent hyper-eutrophication over the 20^th^ century due to increase in nutrient loads from the watershed. Nutrient status was responsible for restructuration of the lake communities across trophic levels (Crosta 1999; Bruel et al. 2018). Air temperature is now driving changes in plankton communities (Bruel et al. 2018).

In a previous study, Cladoceran assemblage was reconstructed at a 2-3 years resolution, from a 74-cm sediment core covering the 1816(±26)—2010 time period (Bruel et al. 2018). Our objective was to evaluate whether the same changepoints could be identified using less samples. In this previous study, the variability in the community was summarized into independent axis using Detrended Component Analysis (Hill and Gauch 1980). Changepoints were then detected on the first component (46% of the total variability) in years 1926, 1946, and 1983. We defined these as the “true” changepoints given they came from an analysis of the complete data. The 1983 changepoint was the largest in magnitude, hence the changepoint we sought to find with our method. We also identified second and third changepoints (Fig. S3).

In line with our simulation approach, we subsampled the full record (74 observations) using the three methods described earlier (random, regular, iterative). These subsamples were from the initial community dataset (Fig. 6a). We reduced the dimensionality of the assemblage-level data to an ordination axis using the same method than in the original study, and detected the changepoint on the first component (univariate vector). In the case of the iterative method, a new sample was added and the ordination was run again (Fig. 6d). For each of the three methods, we examined the error (difference between the true changepoint and the detected changepoint) when using different numbers of samples.

**Figure 4:**
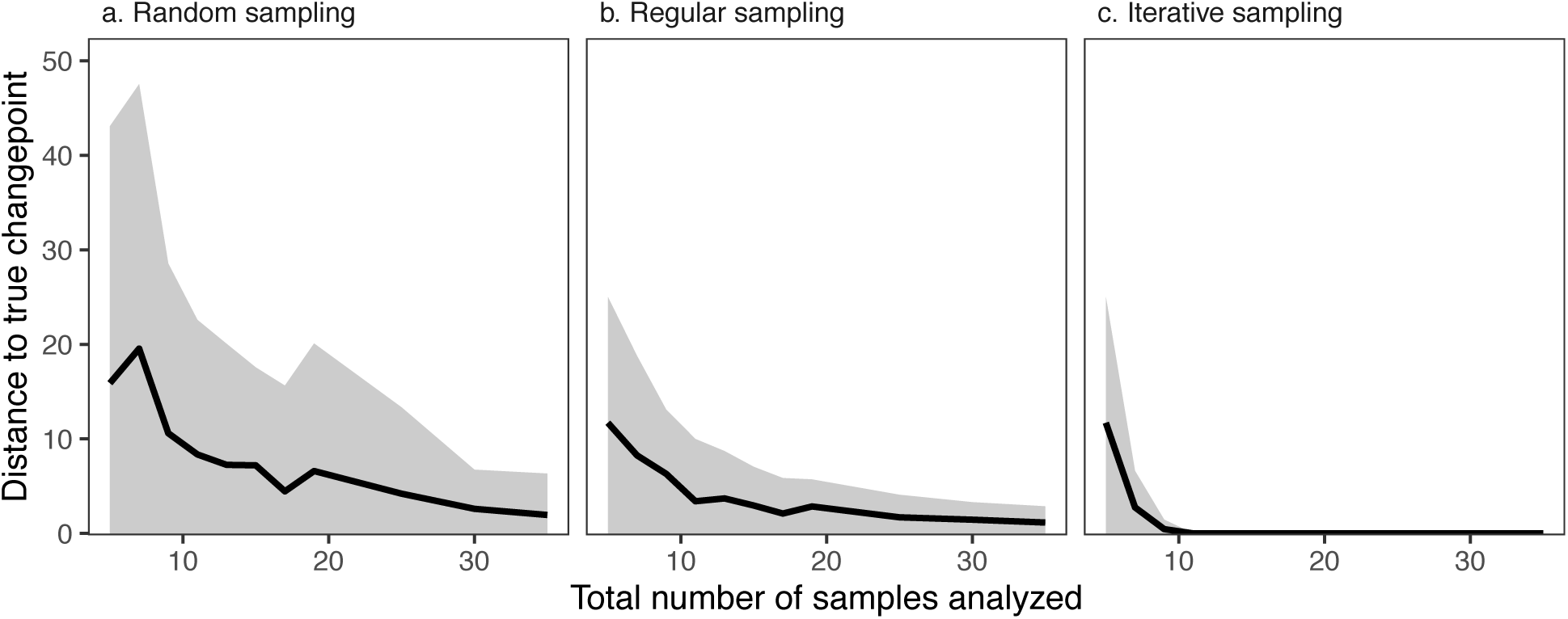
Distance to true changepoint per total number of samples analyzed for simulations, following (a) random sampling, (b) regular sampling, and (c) iterative sampling. Each sample size and approach combination was simulated 50 times with the same parameters as Figure 3. The error bars represent the middle 95% of the simlations.

**Figure 5:**
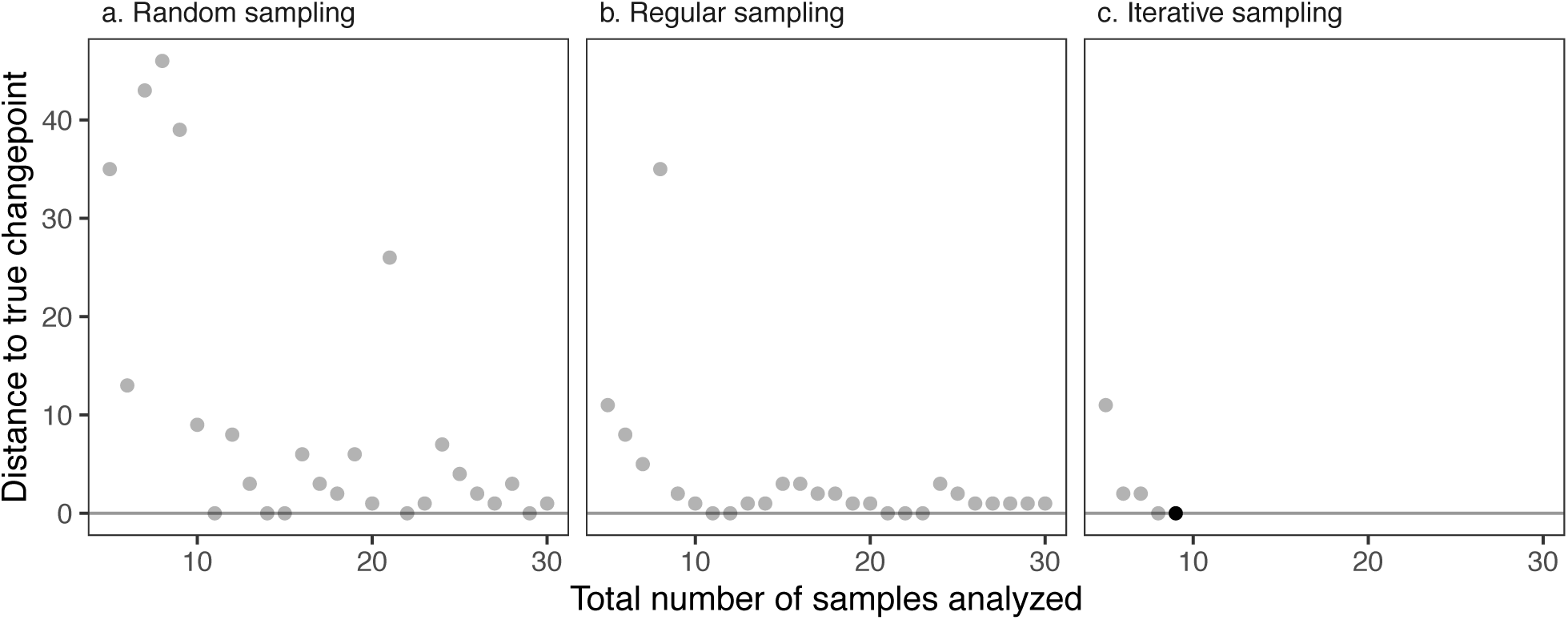
Distance to true changepoint for total number of samples analyzed, following (a) random sampling, (b) regular sampling, and (c) iterative sampling. Total number of samples was set between 5 and 30, out of the 74 initial time series.

**Figure 6:**
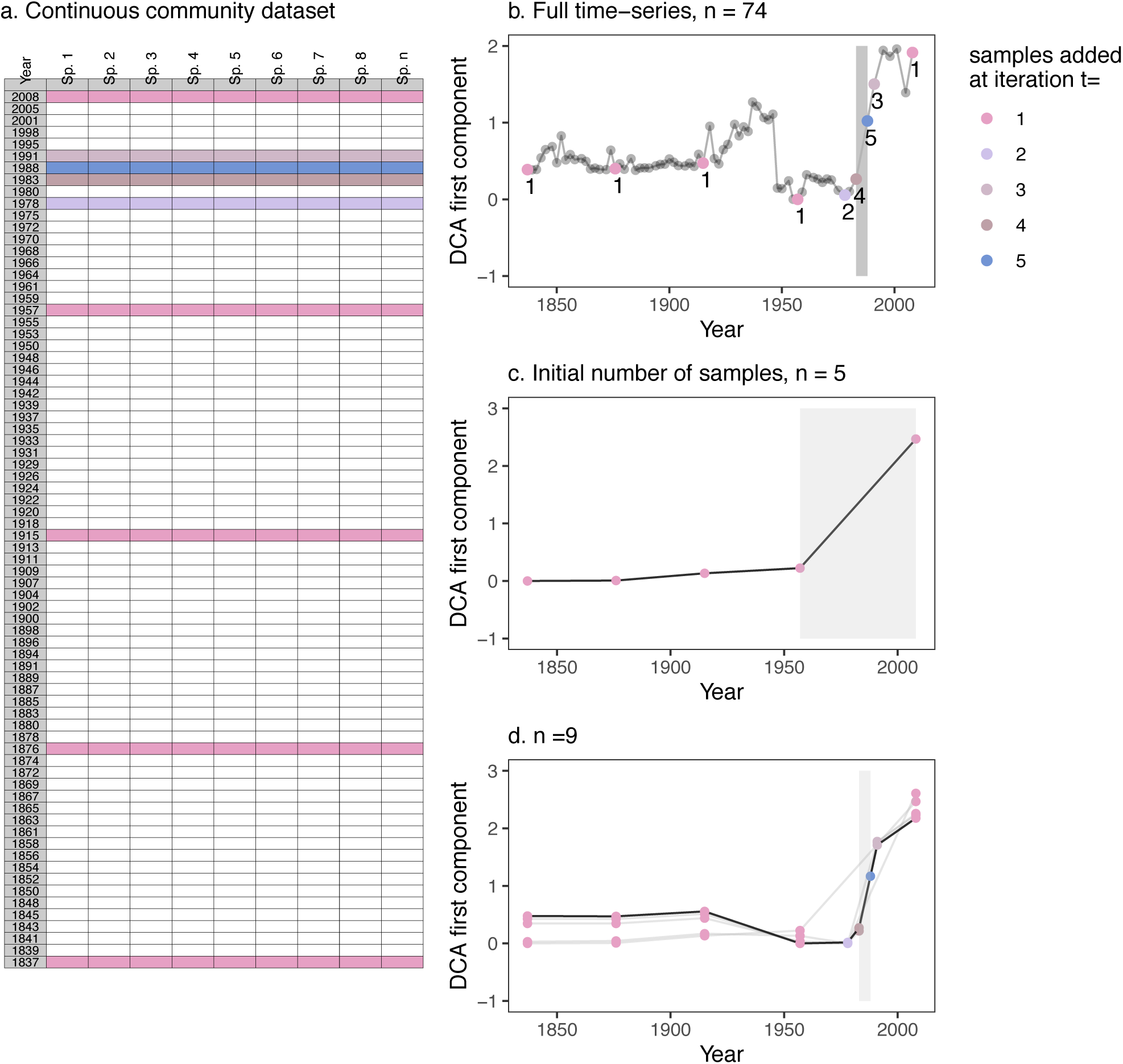
Iterative sampling and impact on ordination. (a) Initial community dataset. Samples are initially sampled at regular interval. The multivariate data is converted to univariate vector by Detrended Component Analysis. Samples were iteratively added, after computation of changepoint at each step. (b) Full time series, on which “true” changepoint was detected. (c) First step of iterative sampling, n = 5. (d) Final step of iterative samples: by adding 4 samples, the true changepoint was detected.

### Case study results

We found that random sampling performed the worst, as changepoint analysis was left to chance (Fig. 5). Regular sampling provided good estimates from 8 samples, but detecting the true changepoint depended on the interval falling close to the true changepoint (i.e., left to chance). Iterative sampling performed the best, as no more than 9 samples were ever necessary to get the true changepoint (Fig. 5c). We show how iterative sampling slightly changes the scores on the first component but not the overall ordination, as more samples are added (Fig. 6)

We also tested how the three methods performed at detecting other changepoints of lower magnitude (as three changepoints were detected in the initial study, Bruel et al. (2018)). Iterative sampling still performed best, especially if an higher number of initial samples was chosen (Fig. S3).

As another application of our approach, we examined the same sediment core data, but examined total abundance as opposed to community composition (Fig. S4). We tested the three subsampling methods, and it took 11 samples to find the “true” changepoint (Fig. S5). The initial 5 subsamples analyzed were the same than the subsamples analyzed to answer the question of the change in community (Fig. 5c).

## Discussion

Due to time or funding limitations, there is often a difference between the number of samples collected and the total number of samples that can later be processed. When the processing time is disproportionately higher than the collection time especially, a subsampling can be done prior to processing. A decision must then be made as to which subsamples to analyze. To address this question in the context of detecting changepoints, we tested three subsampling methods: subsampling random points, regular intervals, and an iterative sampling approach (Fig. 1). We found that the iterative method was systematically better at detecting changes than the two other methods, random subsampling being the least efficient (Figs. 4, 5, S1, S2).

Autocorrelation, variance, and shift size, had an effect on how many samples were needed to detect the shift, but did not change which approach was optimal (Fig. 3).

Multiple subsampling strategies can be chosen (Fig. 1), but only iterative sampling detected the true changepoint with a limited number of samples (Fig. 4c). Analyzing 11% of the sample was enough in most cases to approach the “true” changepoint. Applied to the real case study, the iterative method allowed us to find the main changepoint with only 9 samples analyzed (Fig. 5). The method also worked well to detect other changepoints of lower magnitude (Fig. S3). Bruel et al. (2018) processed one sample at each centimeter in a 74-centimeter sediment core. Each sample took an average of 3 hours to process. We found that using an iterative approach would have eliminated 195 hours of sample processing, or about 24 days, which is just a little over a month of work. This correspond to several thousands of US dollars depending on labor costs.

Our approach goes beyond just paleoecological analyses. Running simulations or using past data to understand the amount of sampling effort required is important in many systems where sample collection or processing is expensive (White 2019; White and Bahlai 2020). The specific sampling techniques can also be compared to determine the optimal strategy in terms of accuracy and cost. Our specific approach applies to situations where more subsamples can be added, or processed, after the dynamics occurred (Zhang and Zhang 2012). It corresponds very well to paleoecological data: samples are taken long after the phenomenon of interest occurred, and allows subsampling at finer or rougher intervals (Wingard, Bernhardt, and Wachnicka 2017). However, both different types of data and different questions than those used here can be addressed with the same approach. Suppose instead that the goal was to detect a change in relative abundance over time with video-based approaches. It is often not practical to watch entire videos, so it can be useful to choose strategic time-points that would address a specific question of interest. Using an interval sampling approach, one could take a fixed number of samples to start. The trend over time from simple linear regression could be taken. Then, samples can be taken at random locations one-by-one and to see which samples have the largest effect on the trend estimates. If a particular sample has a large effect on the trend, then it would be best to choose another nearby sample. Sampling would continue until the trend estimate reached an equilibrium. Thus, the iterative sampling approach is particularly relevant to data sources where additional samples can be taken long after the initial dynamics. These approaches would also be appropriate for environmental samples, such as water or soil, that can be analyzed later or eDNA that can be extracted from previously-collected samples (Bohmann et al. 2014).

Our approach is applicable to a wide range of systems and questions, but it does have limitations. When less resources are needed for sample analysis, as opposed to collection, investigators will likely be able to process every sample, and analyzing all samples to obtain a whole picture may be preferred. We note that if resources need to be saved by collecting less samples in the first place, then regular sampling performs better than random sampling (Figs. 4, S1, S2). Another example where our method is less useful is when addressing questions that require a continuous time series, or at least a regular sampling interval. For example, continuous, high-resolution subsampling of a time-series is generally required to detect critical slowing-down or early warning of shifts (Frossard et al. 2015; Doncaster et al. 2016).

However, recent work suggest that combining indicators (in the specific study, trait dynamics and abundance-based early warning signals) allows forecasting population collapses even with at lower resolution and time-series length (Arkilanian et al. 2020). Critical slowing down does not necessarily result in a shift, and a shift can occur without critical slowing down (Spears et al. 2017). Signs of critical slowing downs are important to understand and recognize because they provide potential early warnings (Doncaster et al. 2016), but in terms of management, knowing the timing of a shift can have larger implications in addressing the underlying driver. Thus, selecting a set number of samples or specific approach may also limit what future questions can be asked.

## Conclusions

Analyzing a subsample of a time series as opposed to the whole time series will inevitably leads to a lesser understanding of the phenomenon observed (White 2019). We show here that an informed subsampling can still allow detection of critical information, such as a changepoint in a time series. Monitoring programs have to be able to address our questions of interest with sufficient statistical power. In addition, optimizing sampling efforts is valuable given the high costs of many monitoring programs (Caughlan 2001; Bennett et al. 2014). Thus, costs of monitoring have to weighed against the value gained from monitoring—a value of information approach (Lovett et al. 2007; Bennett et al. 2018). Monitoring programs should try to anticipate the potential questions of tomorrow, and reducing the data collected, or analyzed, must be done with the best foresight possible on how these data may be necessary to manage ecosystems in the future. If only a subsample of the samples can be analyzed, it may be better to choose samples carefully as opposed to random or regular sampling. This can improve the accuracy of the results and reduce costs overall.

## Code availability

Data and code for all the figures can be found at https://github.com/rosalieb/temporal-sampling.

