## Supplementary Material for "Sampling requirements and approaches to detect ecosystem shifts"

### Contents

**Additional methods** **S2**

**Minimum time for other sampling approaches** **S3**

Random sampling . . . . . . . . . . . . . . . . . . . . . . . . . . . . . . . . . S3

Iterative sampling . . . . . . . . . . . . . . . . . . . . . . . . . . . . . . . . S4

**Detecting further changepoints** **S4**

**Case study: Changepoint detection of abundance time series** **S7**

**References** **S9**

Data and code for all the figures can be found at (https://github.com/rosalieb/temporal-sampling). All the analyses were run in R (R Core Team 2019).

### Additional methods

We first detect the "true" changepoint of a full length time series with the function *e.divisive* in the R package *ecp* (James, Zhang, and Matteson 2019). We focus on the changepoint with the largest magnitude, although this package allows detection of further changepoints as well (Fig. S3). We then subsample the full time series (White and Bahlai 2020) with different numbers of subsamples and different sampling approaches. For each sampling approach, we wrote a custom function:

- sample_random(): sample first and last sample, in addition of n-2 random samples in between (n = maximum number of sample, chosen by the user),

- sample_regular(): sample first and last sample, in addition of n-2 evenly spaced samples in between (n = maximum number of samples, chosen by the user), and

- sample_iterative(): sample first and last sample as well as 3 evenly spaced samples in between (3 is the default but can be modified), in order to initiate the changepoint detection. Upon detection of the changepoint on this 5 samples time series, a new sample is added between the changepoint and the previous sample, to narrow down the real changepoint. Detection of changepoint and addition of sample is repeated, until it was narrowed down to two consecutive samples, or n (maximum number of samples, chosen by the user) was reached, whichever comes first.

The subsampled time series are then compared to the full time series to assess the effectiveness of each subsampling approach.

Each function works with two types of input: vector and matrix. If the input is a matrix, the user must change the default argument *is_vector* to *FALSE*. The matrix is then transformed to independent vectors using Detrended Correspondence Analysis (Hill and Gauch 1980), and a single axis on which changepoint analysis is run is chosen with the argument *DCA_axis* (default to 1, for first component). Component scores are returned by the function. A user can edit the function to use another ordination method (e.g., principal component analysis).

### Minimum time for other sampling approaches

#### Random sampling

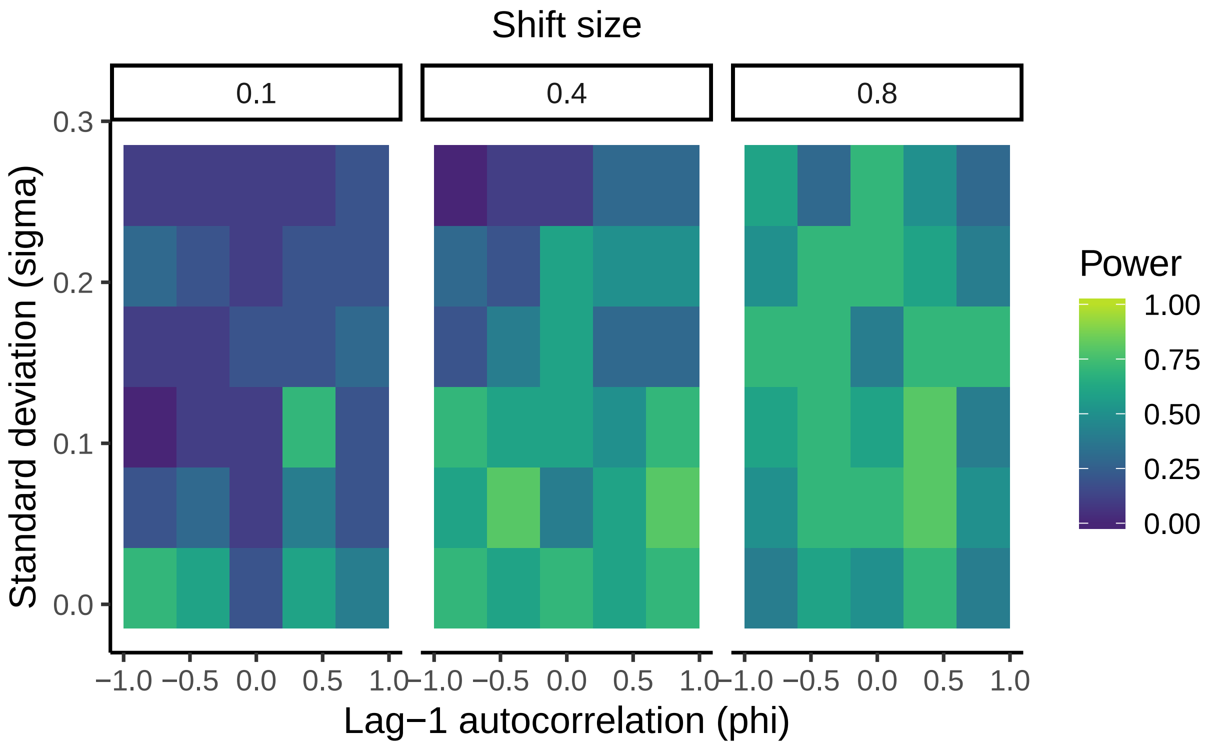


Figure S1: Random sampling statistical power (fraction of 100 simulations which detected a changepoint within five time points of the true changepoint) for different levels of standard deviation ($\sigma$), lag-1 autocorrelation ($\phi$), and shift size ($\delta$). For each parameter combination, 20 samples were used. An increase in samples would increase the statistical power across this graph.

##### Iterative sampling

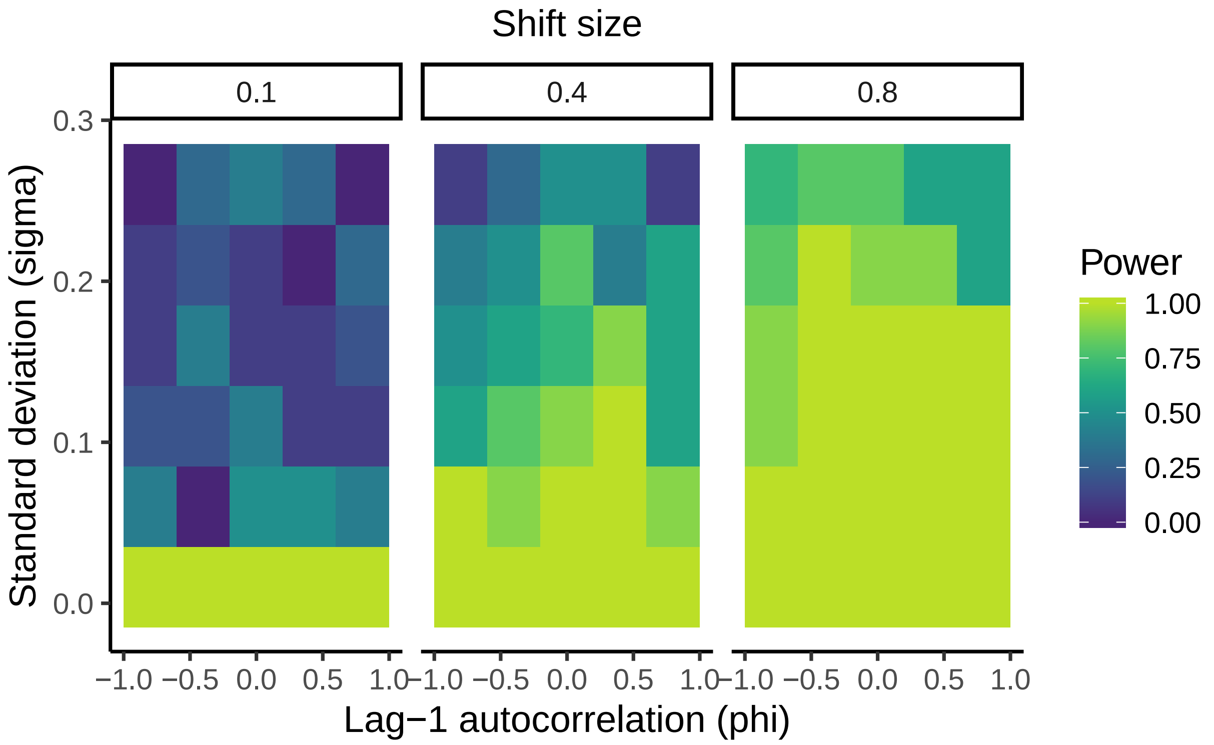


Figure S2: Iterative sampling statistical power (fraction of 100 simulations which detected a changepoint within five time points of the true changepoint) for different levels of standard deviation ($\sigma$), lag-1 autocorrelation ($\phi$), and shift size ($\delta$). For each parameter combination, 20 samples were used. An increase in samples would increase the statistical power across this graph.

#### Detecting further changepoints

The initial study found 3 changepoints on the main axis, in 1926/1928, 1946/1948 and 1983/1988 (Bruel et al. 2018). We focused our analysis on the main changepoint, but the code also allows to target further changepoints, using the argument *c*. *c* is a numeric value indicating the maximum number of changepoints to find (Fig. S3).

The iterative method requires less subsamples overall to find the real changepoints (Fig. S3, 3rd column). Note that if the initial number of samples (controlled with the argument *n2* in the code) is at the default 5, the code struggles to rank appropriately the changepoints, resulting in a changepoint being found, but not the overall "true" second (e.g., Fig. S3b, 3rd column), or "true" third. For a time-series, we recommend users examine the p-values returned by the *e.divisive* function, embedded in the function *sample_iterative* output. If the first few changepoints have close, low p-values, it may be useful to add another subsample further away from the presumed changepoint. This avoids the iterative algorithm mistakenly converging on the wrong changepoint. An alternative is to initiate the subsampling with an higher number of samples, e.g., 7 subsamples.

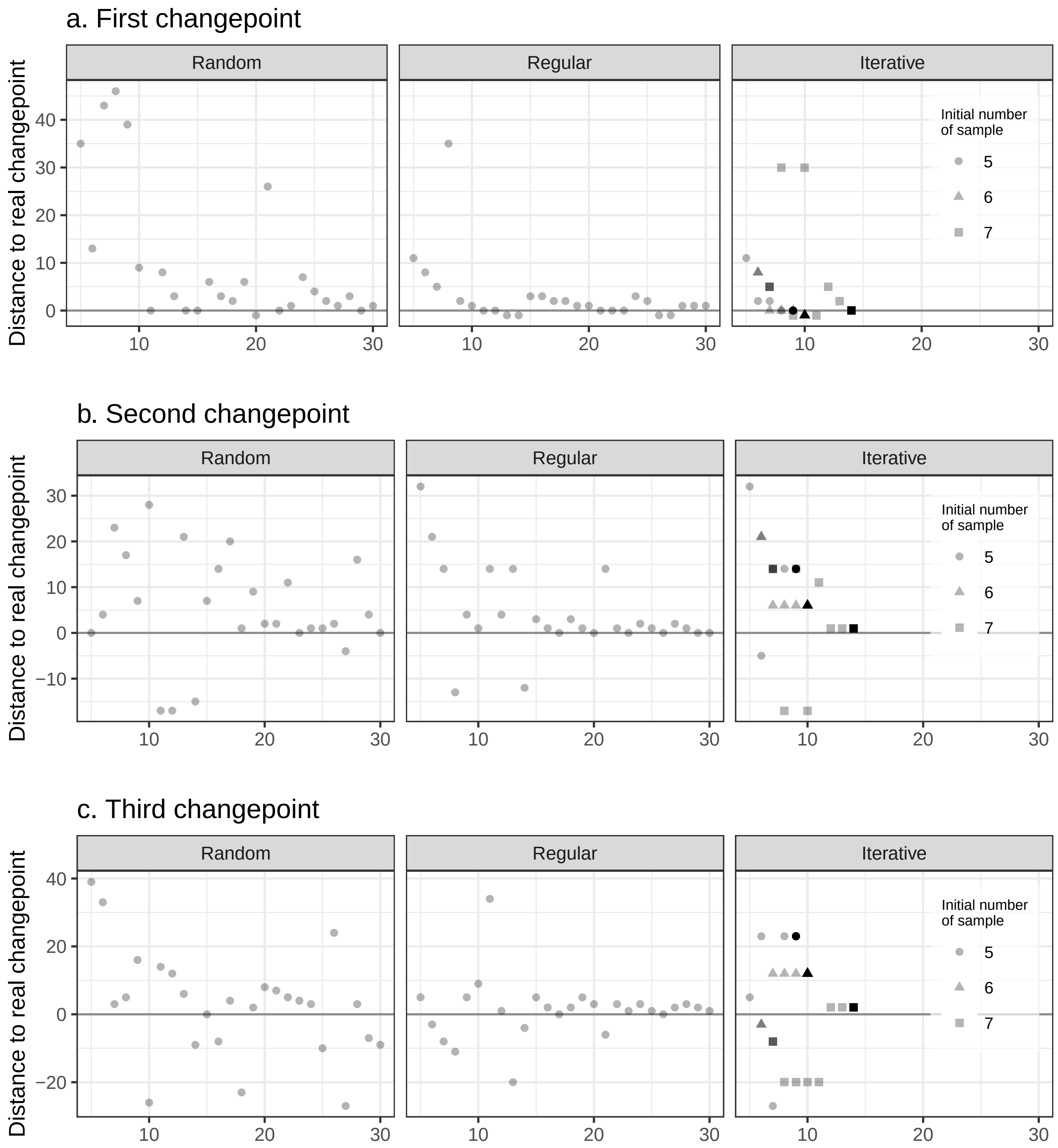


Figure S3: Distance to true (a) first changepoint (largest magnitude), (b) second changepoint, and (c) third changepoint for total number of samples analyzed, following random sampling, regular sampling, and iterative sampling. Total number of samples was set between 5 and 30, out of the 74 initial time series.

### Case study: Changepoint detection of abundance time series

As another example of our approach, we we re-used the community dataset from Lake Varese, but this time, looked at the total number of individuals per gram of sediment in each subsamples. Thus, this is a univariate time series as opposed to the community-level data used in the main text. The full time series show a changepoint in 1946/1948. We used the three changepoint method to estimate changepoint on the vector, and found that the iterative method, with an initial number of 5 regularly-spaced subsamples, performed best (Fig. S5).

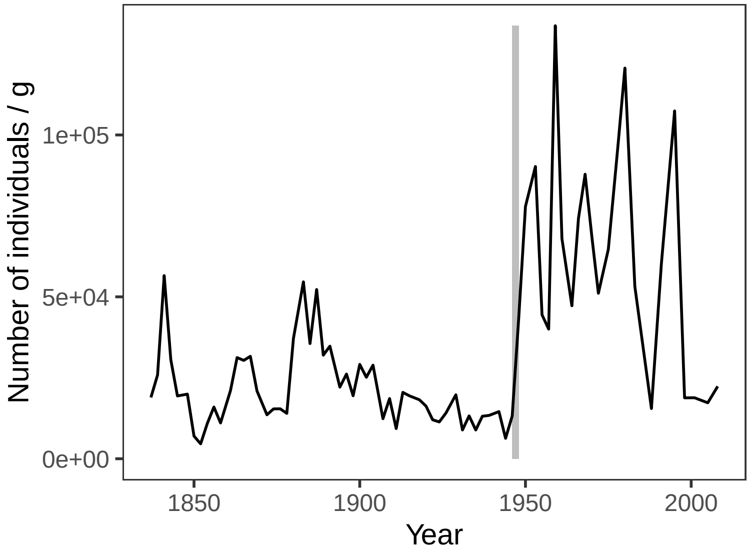


Figure S4: Number of individuals per gram of sediment, infered from Cladocera remains for the 74 subsamples of the Lake Varese sediment core. Vertical grey band indicates the “true” changepoint.

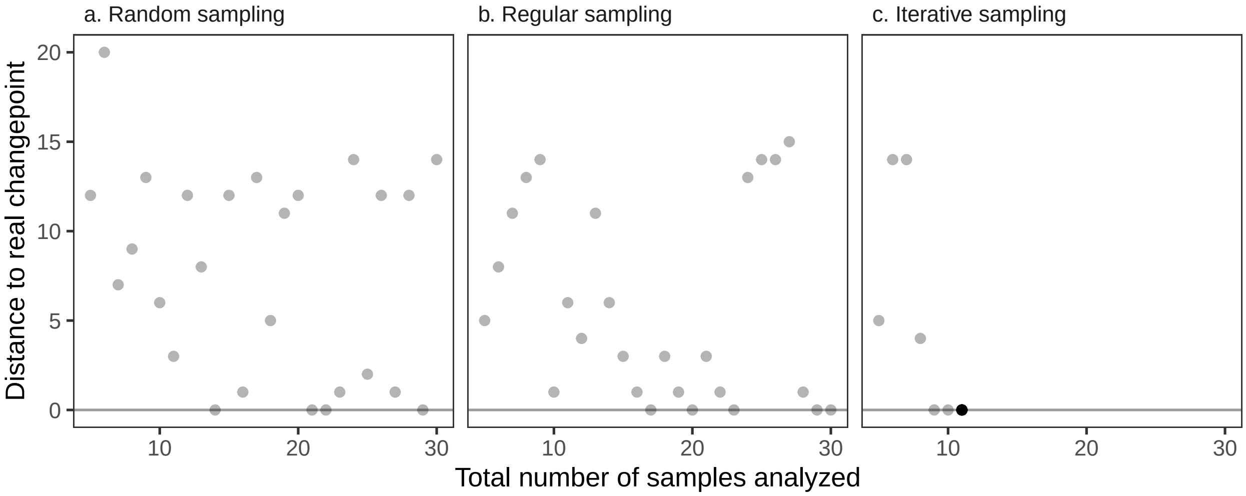


Figure S5: Distance to real changepoint in biomass per total number of samples analyzed, following (a) random sampling, (b) regular sampling, and (c) iterative sampling. Total number of samples was set between 5 and 30, out of the 74 initial time series.

James, Nicholas A., Wenyu Zhang, and David S. Matteson. 2019. "Ecp: Non-Parametric Multiple Change-Point Analysis of Multivariate Data."

R Core Team. 2019. "A Language and Environment for Statistical Computing." Vienna, Austria: R Foundation for Statistical Computing.

White, Easton R, and Christie A. Bahlai. 2020. "Experimenting with the Past to Improve Environmental Monitoring Programs." Preprint. EcoEvoRxiv. https://doi.org/10.32942/osf.io/cz5va.
